# Highly tamoxifen-inducible principal-cell-specific Cre mice with complete fidelity in cell specificity and no leakiness

**DOI:** 10.1101/232108

**Authors:** Lihe Chen, Chao Gao, Long Zhang, Ye Zhang, Enuo Chen, Wenzheng Zhang

## Abstract

An ideal inducible system should be cell-specific and have absolute no background recombination without induction (i.e. no leakiness), a high recombination rate after induction, and complete fidelity in cell specificity (i.e. restricted recombination exclusively in cells where the driver gene is expressed). However, such an ideal mouse model remains unavailable for collecting duct research. Here, we report a mouse model that meets these criteria. In this model, a cassette expressing *ER^T2^CreER^T2^ (ECE)* is inserted at the ATG of the endogenous *Aqp2* locus to disrupt Aqp2 function and to express ECE under the control of the *Aqp2* promoter. The resulting allele is named *Aqp2^ECE^.* There was no indication of a significant impact of disruption of a copy of *Aqp2* on renal function and blood pressure control in adult *Aqp2^ECE/+^* heterozygotes. Without tamoxifen, *Aqp2^ECE^* did not activate a Cre-dependent red fluorescence protein (RFP) reporter in adult kidneys. A single injection of tamoxifen (2 mg) to adult mice enables *Aqp2^ECE^* to induce robust RFP expression in the whole kidney 24h post injection, with the highest recombination efficiency of 95% in the inner medulla. All RFP-labeled cells expressed principal cell markers (Aqp2 & Aqp3), but not intercalated cell markers (V-ATPase B1B2, and carbonic anhydrase II). Hence, *Aqp2^ECE^* confers principal cell-specific tamoxifen-inducible recombination with absolute no leakiness, high inducibility, and complete fidelity in cell specificity, which should be an important tool for temporospatial control of target genes in the principal cells and for Aqp2+ lineage tracing in adult mice.

## Introduction

Tamoxifen-dependent CreER recombinases have been developed for disruption or activation of target genes at will in a temporospatially-controlled manner (4). They consist of Cre fused to a mutated ligand-binding domain (LBD) of the estrogen receptor (ER). The mutated LBD binds with a high affinity the synthetic ER ligand tamoxifen and its active metabolite 4-hydroxytamoxifen (4OHT), but not the endogenous estrogens. In the absence of the ligand, the CreER fusion protein remains inactive in the cytoplasm. In the presence of tamoxifen, the CreER fusion binds tamoxifen and moves into the nucleus where it recombines its loxP-flanked DNA substrate. Currently, the most successful CreER version is CreER^T2^, which harbors the human ER LBD with a G400V/M543A/L544A triple mutation. By breeding CreER^T2^ driver with mice containing floxed essential exon(s) of target genes, gene targeting can be temporally regulated. To achieve spatial control, the expression of *CreER^T2^* is placed under the control of the various tissue/cell-specific promoters.

The kidney contains many functionally and structurally different cells. Several segment-specific *CreER^T2^* transgenic or knock-in mouse lines have been reported, using promoters of *Gamma glutamyl transpeptidase (GGT)* (7), *Podocin* (23), *Cited1* (3), *Lgr5* (2), *Six2* (9), *Zfyve27* (16), *Foxd1* (8), and *Ksp-cadherin* (11, 17). In addition, transgenic or knock-in mice expressing *Tet-off-eGFPCre* and *eGFPCreER^T2^* driven by the promoters of *Foxd1* (8), *Six2* (9), and *SLC34a1* (10), respectively, are also available. Many of these inducible Cre lines are active in multiple segments, particularly in proximal tubules.

In *Aqp2Cre* (15, 20) and *Aqp2CreER^T2^*(21) mice *Aqp2* serves as the driver gene since its promoter drives Cre and CreER^T2^ expression, specifically in connecting tubule/collecting duct (CNT/CD). These models have been used by others (1, 18, 20, 21, 27) and us (25) to create CNT/CD-specific knock out mice. However, the constitutive nature of the *Aqp2Cre* (15, 20) and the high background recombination in the absence of tamoxifen (i.e. leakiness) of the *Aqp2CreER^T2^* (21) do not offer temporal control, limiting their use in studying the role of CNT/CD in pathological conditions of the adult kidney.

Electroporation studies demonstrated that *ER^T2^CreER^T2^ (ECE)* had no leakiness (13). To our knowledge, the tight control of *ECE* has not been strictly tested *in vivo* since *ECE* transgenic or knock-in mice have not been reported. Here, we report a new inducible mouse model in which an *ECE* cassette is inserted into mouse genome at the ATG of the endogenous *Aqp2* locus. The resulting allele is referred to *Aqp2^ECE^.* We demonstrate that *Aqp2^ECE^* has very mild or no effect on renal function, absolute no leaky Cre activity in the absence of tamoxifen, high recombination efficiency upon induction, and specific recombination exclusively in the cells where *Aqp2* expression occurs (i.e. complete fidelity or faithfulness in cell specificity). Our study defines *Aqp2^ECE^* as a new powerful system for evaluating gene function specifically in the principal cells at any time and for identifying Aqp2^+^ progenitor cells in adult kidney. Our *ECE* knock-in strategy can also be applied to develop various tissue/cell-specific *ECE* drivers that may also have high inducibility, complete faithfulness and no leakiness.

## Materials and Methods

### Reagents

Two mouse antibodies specific for <u>carbonic anhydrase II</u> (CAII, sc-48351) and V-ATPase B1 B2 (sc-55544), one rabbit Aqp2 antibody (sc-28629), and two goat antibodies against Aqp2 (sc-9882) and Pendrin (sc-16894) were obtained from Santa Cruz, Dallas, Texas, USA. Rabbit-anti RFP (632496, Clontech, Mountain View, CA, USA), and chicken anti-GFP (ab13970, Abcam, Cambridge, MA, USA) were also used. The secondary antibodies were purchased either from Invitrogen, Carlsbad, California, USA or Jackson ImmunoResearch, West Grove, PA, USA. They were Alexa 647 donkey anti-goat IgG (A21447), Alexa 568 donkey anti-rabbit IgG (A10042), Alexa 488 donkey anti-goat IgG (A11055), Alexa 488 donkey anti-chicken IgG (703-545-155) and Alexa 488 donkey anti-mouse IgG (715-545-150). Tamoxifen (T5648), sunflower oil (S5007) and all primers (Table 1) were ordered from Sigma-Aldrich, St. Louis, Missouri, USA. pCAG-ERT2CreERT2 was obtained from Addgene, Cambridge, Massachusetts, USA. Arg^8^-Vasopressin ELISA kit (ADI-900-017A) was purchased from Enzo, Farmingdale, NY, USA.

**Table 1.**
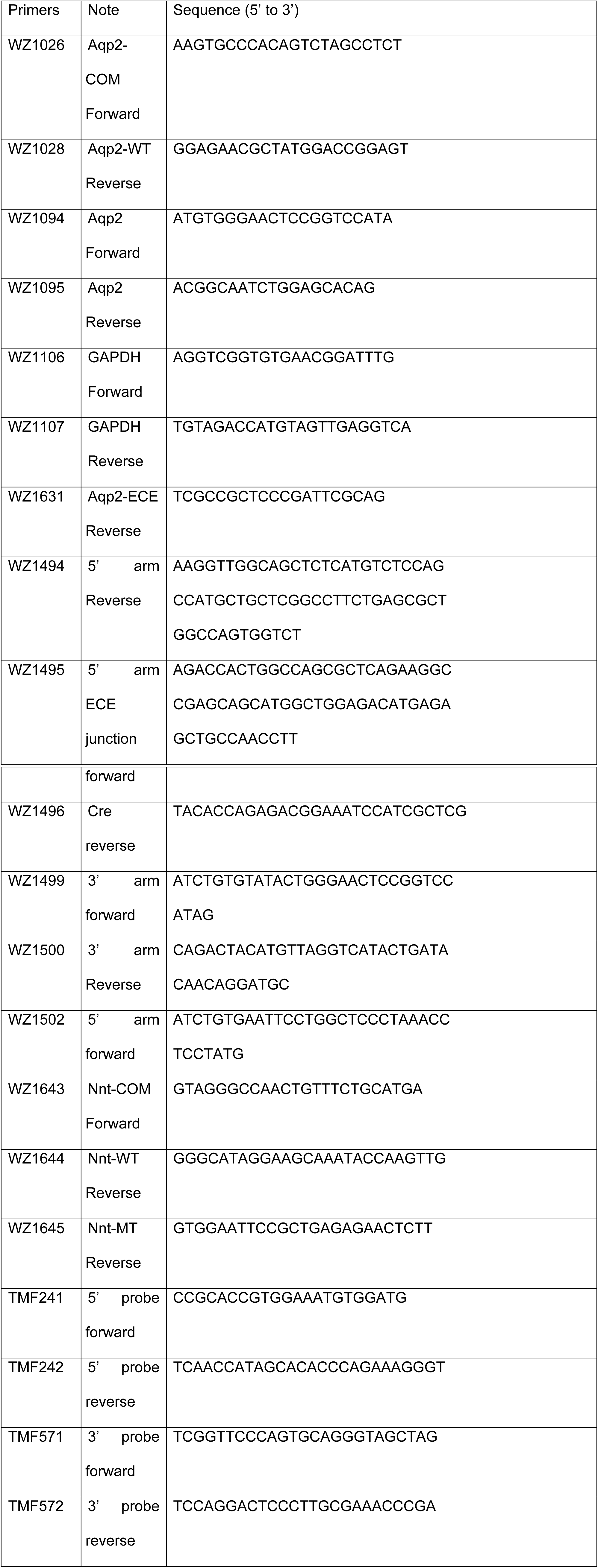
Primer list

### Generation of the targeting vector, ES cells, and chimeras

A 2.5kb PCR fragment of *Aqp2* gene as the 3’ arm was amplified using primers WZ1499/1500 with mouse tail DNA as template. The fragment was cloned into pcDNA 3.1 (+) at PciI and BstZ171 to create p692. A 2.9kb fragment containing *ER^T2^CreER^T2^* was released from pCAG-ERT2CreERT2 and inserted into p692 at EcoRI and NotI to generate p693. The 5’ arm was created through overlapping PCR. The first 4.1kb fragment of *Aqp2* gene was amplified from mouse genomic DNA with primers WZ1494/1502. The second 1.6kb fragment was synthesized with primers WZ1495/1496 using pCAG- ERT2CreERT2 as template. The final 5.7kb 5’ arm obtained via PCR using primers WZ1496/1502 with the two fragments as the template was then cloned into p693 at EcoRI and AgeI to generate the final target vector p694. The authenticity of the whole insert sequence in the final target was confirmed by sequencing.

After linearization with KpnI, which trimmed the 5’ and 3’ arms to 1.8kb and 2.3kb, respectively, the targeting vector was electroporated into C57BL/6N JM8.N4 ES cells. The C57BL/6N background of the ES cells offers a unique advantage for identification of chimeras (see below). A total of 384 G418 resistant clones were picked, duplicated, expanded and total DNA extracted for Southern analyses. Probes to screen for homologous recombination in Southern analyses were selected that were outside of the region of homology used for the arms of the targeting vector. Probes were generated using PCR. The 3’ probe (893bp) was amplified using primers TMF571/572. The 5’ probe (474bp) was amplified using primers TMF241/242. DNA fragments were labeled using α-^32^P-dCTP and hybridized and washed to digested ES cell genomic DNA using standard methods. To detect recombination at the 3’ arm, DNA was digested with BamHI, with a 9.8kb hybridizing band indicating wild type and 12.5kb band allele denoting targeted allele. For the 5’ arm, DNA was digested with EcoRV with band sizes of 9.7kb (WT) and 8.6kb (targeted) (Fig 1A). Chromosome counting was carried out to verify the genome integrity of 6 correctly targeted ES clones. Two of these clones with >80% euploid cells were injected into C57BL/6J blastocysts to produce chimeras. Since the ES cells and the blastocysts are homozygous for the WT and mutant nicotinamide nucleotide transhydrogenase gene (*Nnt^+/+^* and *Nnt^−/−^),* respectively (19), PCR analyses were done to identify the chimeras and to evaluate the relative contribution of the ES cells. The expected products are 576bp for the wild-type allele and 743bp for the mutant allele.

**Fig. 1.**
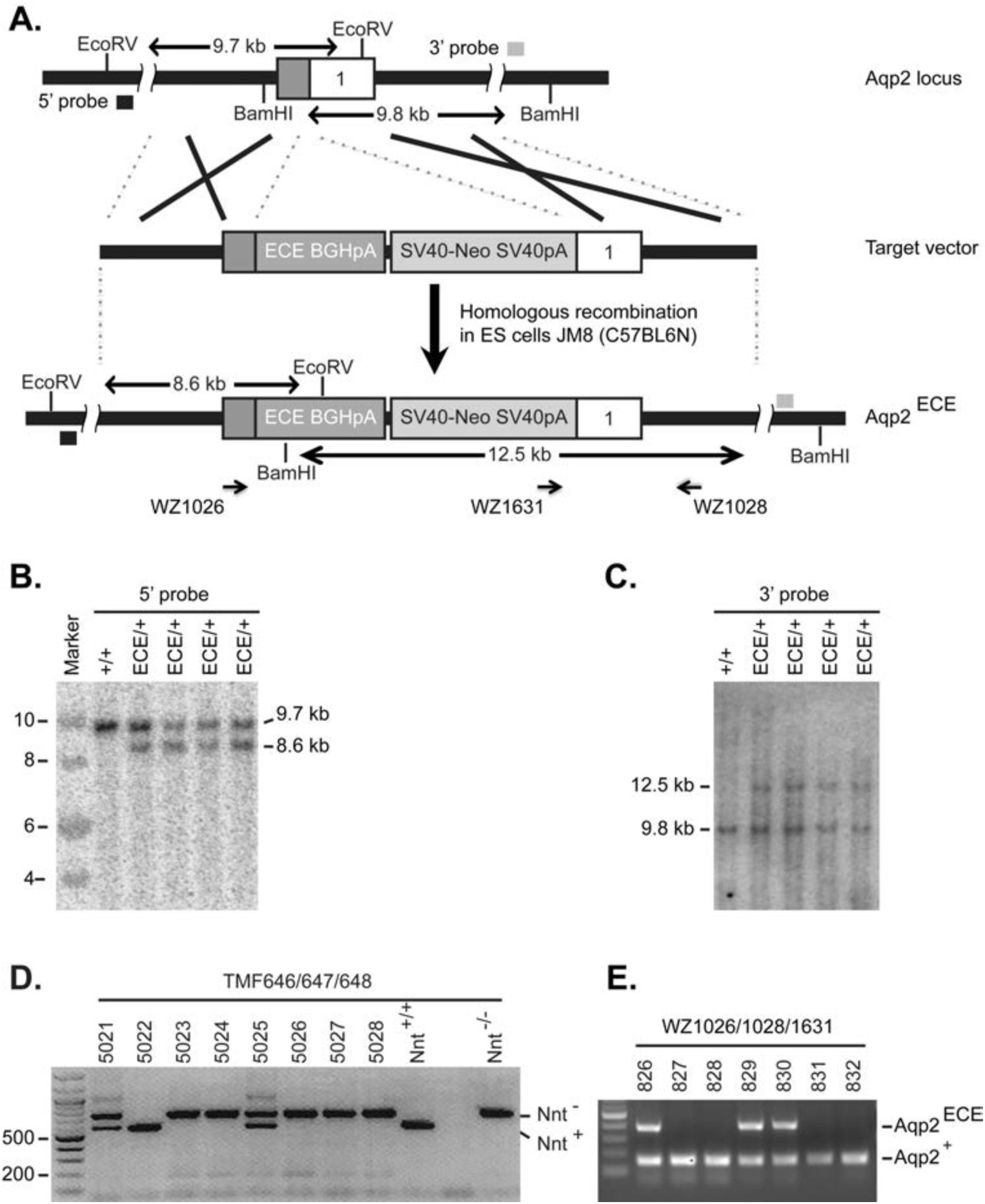
Generation of *Aqp2^ECE/+^* mice. **A.** Schematic illustration of targeting strategy for the *Aqp2^ECE^* knock-in allele. **B-C.** Southern blotting analyses of ES clones confirming the targeted *Aqp2^ECE^* knock-in allele. **D.** PCR-based genotyping of chimeras showing male 5022 with almost 100% contribution of ES cells, as evidenced by the presence of the ES cell-derived WT allele coupled with barely detectable host blastocyst mutant allele of nicotinamide nucleotide transhydrogenase gene (Nnt). **E.** PCR-based genotyping showing that male 5022 had a high rate in germline transmission of *Aqp2^ECE^* allele to offspring.

### Generation and genotyping of *ECE/+ RFP/+* and *ECE/+ RFP/RFP* mice

Male chimeras founders were bred with *R26R-RFP* (Jackson Laboratory stocks 007914 (12)) for germline transmission and for introduction of the Cre-dependent reporter. The resulting mice from successful germline transmission were heterozygous for both the ECE and RFP alleles. They were referred as *ECE/+ RFP/+* mice. *ECE/+ RFP/RFP* mice were heterozygous for the ECE allele, but homozygous for the RFP allele. They were created through crossing *ECE/+ RFP/+* with *R26R-RFP* strain. PCR-based genotyping was conducted to identify the genotypes. Primers used for the WT (150bp) and *Aqp2^ECE^* (380bp) alleles were WZ1026/1028/1631. R26R-RFP allele was genotyped according to the instruction listed in the Jackson Laboratory website. All mice were generated in a highly pure C57BL/6 background, and examined at the age of 2-3 months.

### RT-qPCR and Western blotting analyses

These assays were conducted as we reported (6, 25). Briefly, total RNAs and proteins were isolated from kidneys of un-induced *Aqp2^+/+^* and *Aqp2^ECE/+^* kidneys with free water access (n=3 mice/genotype) and under water deprivation for 24h (n=4 mice/genotype). Each of RNA samples was analyzed via real-time RT-qPCR in triplicates, with GAPDH as internal control. The PCR primers were WZ1094/1095 for Aqp2 (137 bp) and WZ1106/1107 for GAPDH (123 bp). Western blotting analyses were performed using a rabbit anti-Aqp2 antibody (Santa Cruz, sc-28629). The 35-50kd band corresponding to the glycosylated Aqp2 was quantified and normalized to β-actin. In all case, the relative Aqp2 mRNA and protein levels in WT were set to 1.

### Metabolic balance study

*Aqp2^+/+^* and their *Aqp2^ECE/+^* littermates without tamoxifen injection were given a normal Na^+^ diet (LabDiet, 5P76). Mice either had free water access or were under water deprivation for 24h. They were placed in metabolic cages (Tecniplast, 3700M022) with modified feeders to improve food delivery. Urine was obtained via the metabolic cage and used to estimate all urinary parameters. Na^+^ excretion was estimated as the product of urine [Na^+^] times 24h urine volume, and normalized to BW. K+ excretion was calculated similarly. For water deprivation study, residual urine was eliminated at the start of the water deprivation study. Systolic and diastolic blood pressure with the CODA tail-cuff blood pressure system (Kent Scientific, Torrington, CT, USA) was measured as we previously reported (6, 25, 29). Each mouse was received at least one cycle of measurement containing 20-30 individual readings for each parameter each day. To minimize circadian effects and daily variation, water intake, food intake, urine collection, and blood presure measurements were performed around 3 pm each day for three running days. For each mouse, data from multiple days were pooled to calculate the final average to represent that mouse and counted as 1 (n=1).

### Blood and urine measurements

Measurement of blood and urine parameters were conducted as we reported before (6, 25). Briefly, urine [Na^+^] and [K^+^] were measured with a flame photometer (PFP7, Jenway, UK). Urine osmolarity was recorded with a vapor pressure (Wescor Vapro Vapor Pressure Osmometer 5520, Scimetrics, Houston, TX, USA). Urine [AVP] was determined using a ELISA kit as reported (28). Blood was analyzed using VetScan i-STAT (ABAXIS, Union City, CA, USA) according to the manufacturer’s instruction. Blood [glucose] was also measured via a tail snip with a normal glucose meter.

All animal studies were approved by The Institutional Animal Care and Use Committee, Albany Medical College.

### Induction with tamoxifen

Tamoxifen was dissolved in sunflower oil by sonication at 10 mg/ml, aliquoted and stored at -80 °C before use. To minimize the change in tamoxifen activity due to repeated freezing and thawing, each aliquot was used only once. Adult *ECE/+ RFP/+* and *ECE/+ RFP/RFP* mice were given a single intraperitoneal injection of 2 or 4 mg tamoxifen or an equal volume of sunflower oil. Mice were sacrificed for tissue harvest 24h or 2 months post injection.

### Immunofluorescence and confocal microscopy studies

Immunofluorescence staining and analyses of epifluorescence and confocal images were conducted as we reported (24-26), with one exception. Confocal images were taken using a Zeiss LSM 880 NLO confocal microscope with Airyscan at The Albany Medical Center Imaging Center. At least ten fields were taken from the whole kidney in each of *ECE/+ RFP/+* and *ECE/+ RFP/RFP* mice. Cell counting was restricted to tubules consisting of at least one Aqp2^+^ cell or at least one RFP+ cell. DAPI staining was added to aid cell counting. Recombination efficiency was defined as the number of RFP+Aqp2^+^ cells divided by the sum of the number of RFP^+^Aqp2^+^ cells and the number of RFP^−^Aqp2^+^ cells.

### Statistical analyses

Quantitative data were presented as mean±SD. Student t-test was conducted to determine the statistical significance, which was set at P<0.05.

## Results

### Generation of a new inducible *Aqp2^ECE/+^* mouse model

A PAC transgene *(Aqp2CreER^T2^)* with *CreER^T2^* inserted into the *Aqp2* locus at the position of the *Aqp2* initiation codon caused tamoxifen-independent recombination in mice (21), indicating leakiness. Similarly, Cre fusions (*CreER^T2^, ER^T2^Cre,* and *FlpeER^T2^*) caused high background recombination without 4OHT when temporally expressed from electroporated plasmids (13). In contrast, *ER^T2^CreER^T2^ (ECE)* tested in parallel was 4OHT-dependent (13). To create a principle-cell-specific, tightly controlled inducible system, we intended to create an *Aqp2^ECE^* (or *ECE* in short) knock-in allele. To this end, a targeting vector with an *ECE* cassette inserted into the *Aqp2* locus at the position of the *Aqp2* ATG was constructed (Fig. 1A and (5)), and electroporated into C57BL/6N-derived ES cells to knock-in *ECE* at the endogenous *Aqp2* locus and to completely disrupt the function of *Aqp2.* Southern analysis revealed an extremely efficient targeting frequency of > 90% (Fig 1B,C and data not shown). Injection of two correctly targeted ES clones yielded 4 male chimeras. As shown in Fig 1D, male chimera 5022 had almost 100% contribution from the ES cells as indicated by the presence of ES cell-derived WT *Nnt* band coupled with undetectable mutant *Nnt* band. Consistently, it exhibited a high rate in germline transfer of the *ECE* allele to its offspring (Fig. 1E). Two of the other three male founders with 50% to 90% ES contribution also showed germline transmission. The resulting heterozygotes were termed *Aqp2^ECE/+^* or *ECE/+.* Since all *ECE/+* mice derived from the three male founders did not show a particular phenotype and were healthy, viable, and fertile, they were characterized further.

### *ECE/+* mice apparently have normal renal function and blood pressure

To determine if disruption of one copy of *Aqp2* by the *ECE* insertion causes a significant effect on Aqp2 mRNA expression, renal function and BP control, we performed real-time RT-qPCR, Western blotting, metabolic analyses, and BP measurements. In these experiments, uninduced adult mice was given a normal Na^+^ diet either with free access to water or under water deprivation for 24h. Aqp2 mRNA and protein levels were significantly decreased in *ECE/+* vs. WT (n=3-4 mice/genotype, Fig. 2A-C) when they had free access to water. However, such differences were not observed when they were challenged with the water deprivation (n=4 mice/genotype) (Fig. 2D-F). Analyses of 10-16 WT and 11-22 *ECE/+* mice revealed no significant difference in body weight, water intake, food intake, urine parameters (urine volume, osmolarity, [Na^+^], [K^+^], excretion of Na^+^ and K^+^, and [AVP]), blood parameters ([Na^+^], [K^+^], [Cl^−^], [TCO2], [BUN], [HCO_3-_], [BEecf], [AnGap], [Hb], [PCO_2_], [Hct], pH, and [Glucose]), and diastolic and systolic pressure between the two genotypes (Fig. 3A-Z). It should be noted that [glucose] measured with iSTAT was substantially higher than [glucose] measured via a tail snip with a normal glucose meter for both genotypes (Fig. 3W & X).

**Fig. 2.**
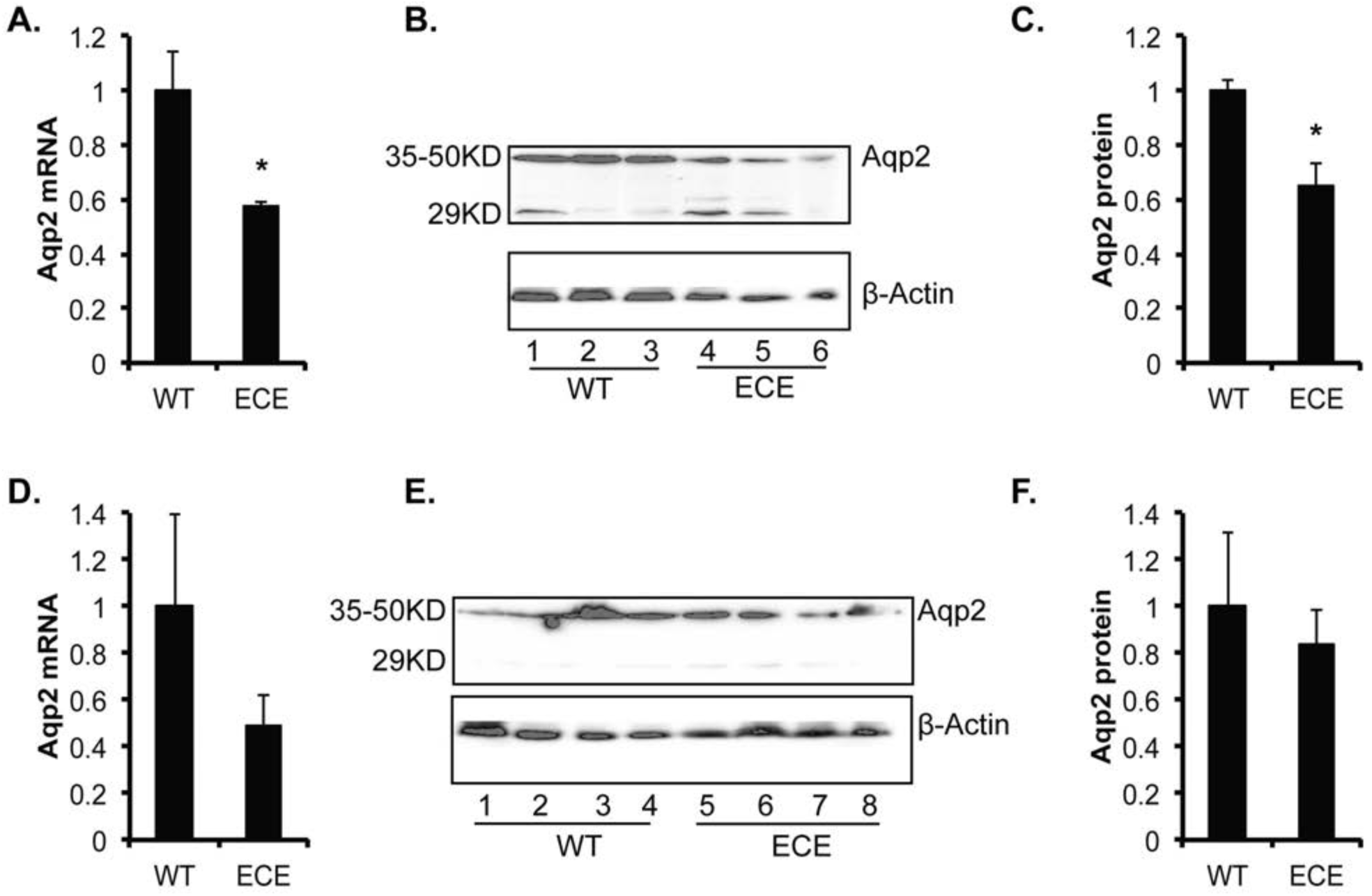
Analyses of Aqp2 expression. Total RNAs and proteins were isolated from kidneys of un-induced *Aqp2^+/+^* and *Aqp2^ECE/+^* kidneys with free water access (n=3 mice/genotype, ***A-C***) and under water deprivation for 24h (n=4 mice/genotype, ***D-F***). Each of RNA samples was analyzed via realtime RT-qPCR in triplicates, using GAPDH as internal control. For Western blotting analyses, the rabbit Aqp2 (sc-28629) was used. The 35-50kd band corresponding to the glycosylated Aqp2 was quantified, normalized to β-actin and presented in ***C*** and ***F***, respectively. In all case, the relative Aqp2 mRNA and protein levels in WT were set to 1. *: P<0.05 vs. WT.

**Fig. 3.**
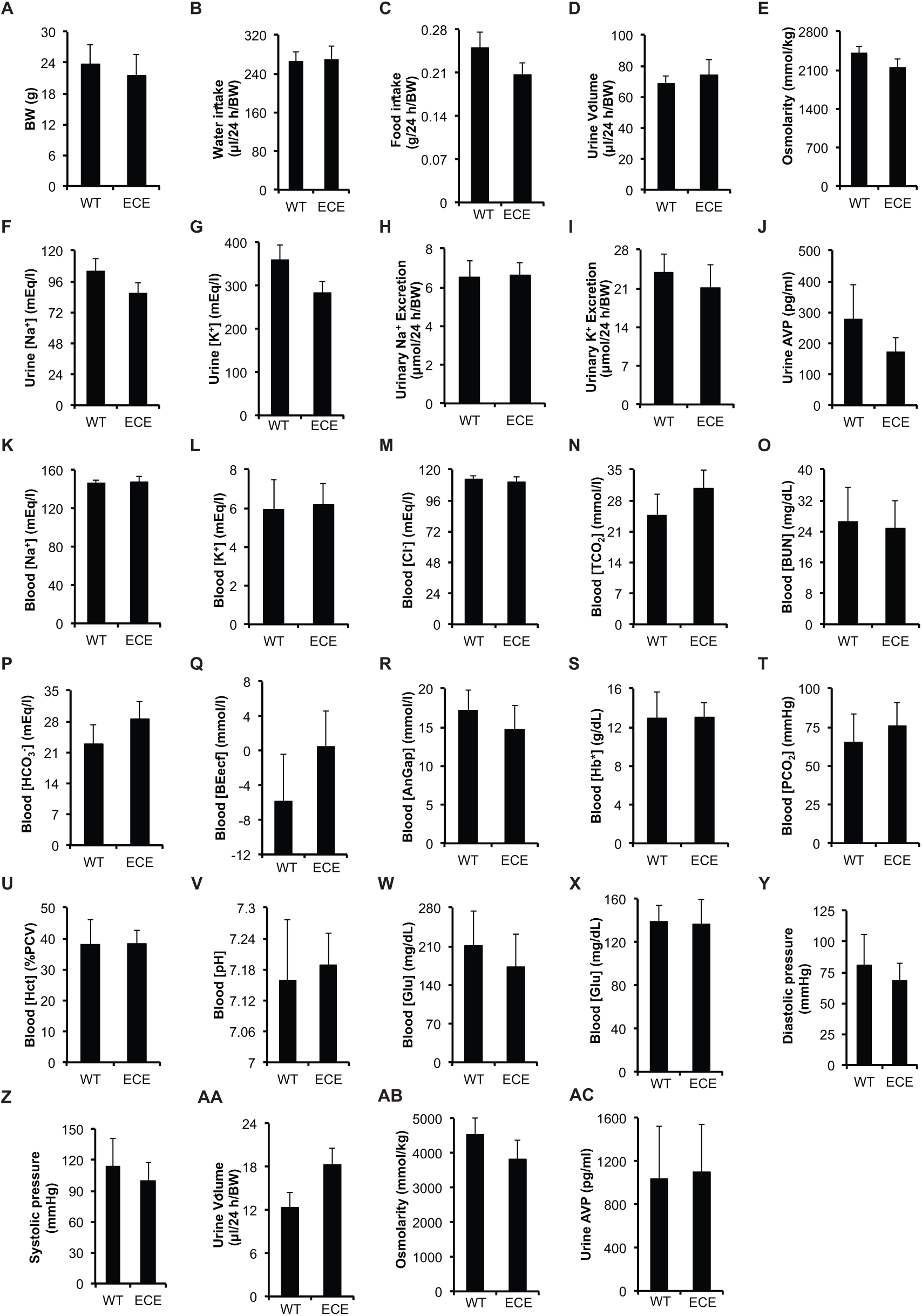
*ECE/+* mice apparently have normal renal function and blood pressure. Uninduced adult WT (n=10-16) and *ECE/+* mice (ECE) (n=11-22) were given a normal Na^+^ diet either with free water access (***A-Z***) or were under water deprivation for 24h (***AA-AC***) were analyzed for the parameters as indicated. K-W were done with iSTAT. X was measured via a tail snip with a normal glucose meter. The significance of the difference at P<0.05 in each of the parameters between the two genotypes was not reached.

ECE/+ vs. WT mice were also indistinguishable in the urine volume, osmolarity and [AVP] after 24h water deprivation (Fig. 3AA-AC). Hence there is no indication of a significant effect of inactivation of one copy of *Aqp2* by the *ECE* knock-in on renal function and BP maintentance under the conditions tested.

### *Aqp2^ECE^* possesses absolute no leaky activity

*ECE* had no detectable recombination activity without 4OHT in electroporation studies (13). However, to our knowledge, the tight control of *ECE* has not been rigorously evaluated when it is permanently expressed *in vivo* since no *ECE* transgenic or knock-in animals have been reported. Accordingly, we bred each of the three male chimera founders with R26R^tdTomato/+^ (referred as RFP) reporter (Jackson Laboratory stock 007914). RFP is expressed after removal of the floxed stop cassette through Cre-mediated recombination. Immunofluorescence staining with the anti-RFP and anti-Aqp2 antibodies verified the lack of RFP expression in the whole kidney in oil-injected two-month old double heterozygotes *ECE/+ RFP/+* and *ECE/+ RFP/RFP* mice that were heterozygous for *ECE,* but homozygous for *RFP* (Fig. 4 and data not shown). Therefore, our data demonstrate absolutely no “leaky” activity of *ECE* in the absence of tamoxifen induction. In addition, our data also indicate that *ECE* transmission can be carried out through males. There are two possibilities. The male germline activity of the *Aqp2* promoter (15) did not cause ECE-mediated background recombination in the sperms. Alternatively, *Aqp2* promoter is inactive in the male germline. The latter is consistent with undetectable Aqp2 and RFP expression in the testis in the induced *ECE/+* mice (see Fig. 10B).

**Fig. 4.**
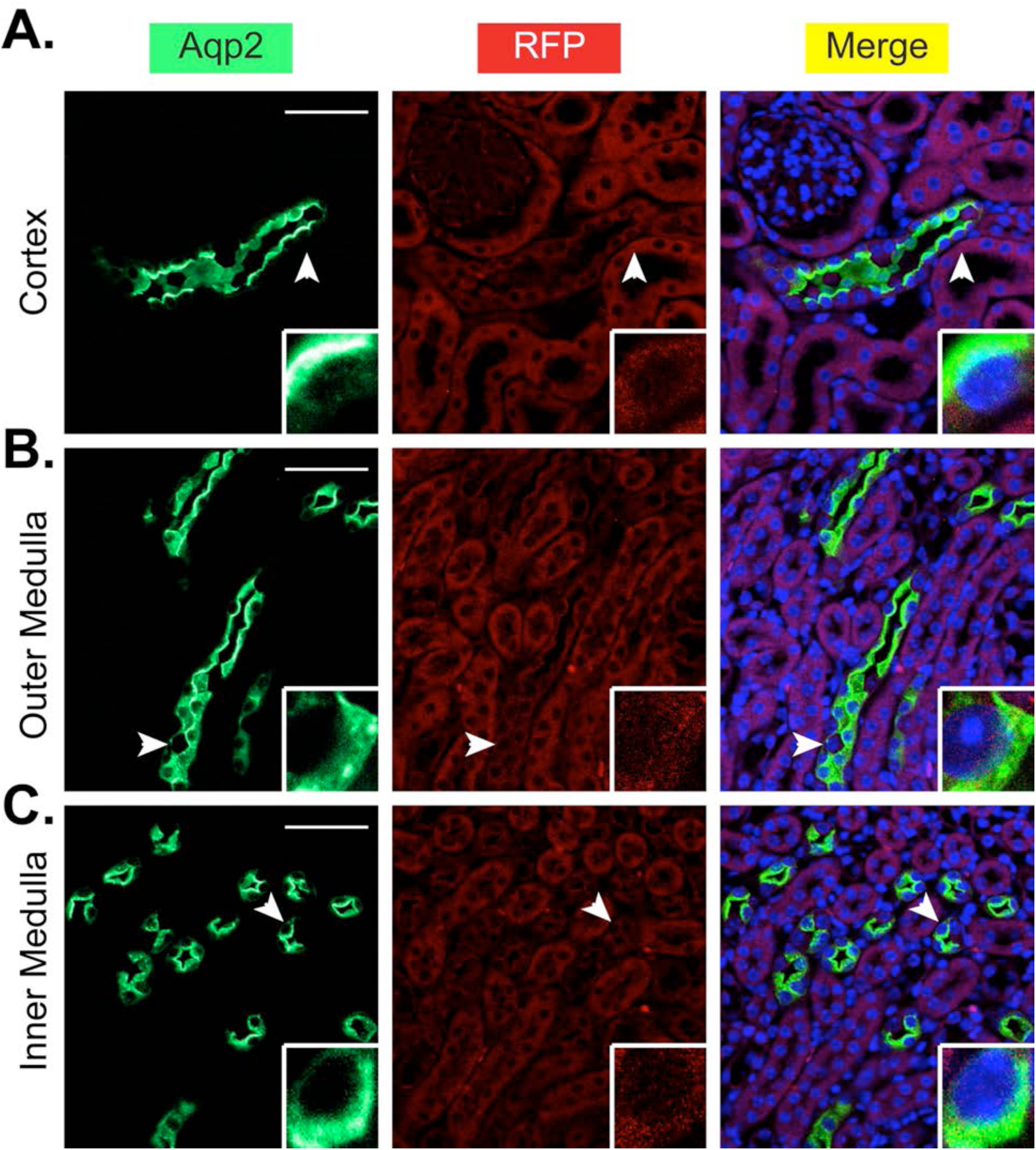
*Aqp2*^ECE^possesses absolute no leaky activity. *A-C*. Immunofluorescence staining for Aqp2 (green) and RFP (red) showing no RFP expression in the whole kidney of an oil-injected adult *ECE/+ RFP/RFP* mouse. Cells indicated by arrowheads were magnified in the inserts. Scale bar: 50 μm and 8.3 μm for insert.

**Fig. 10.**
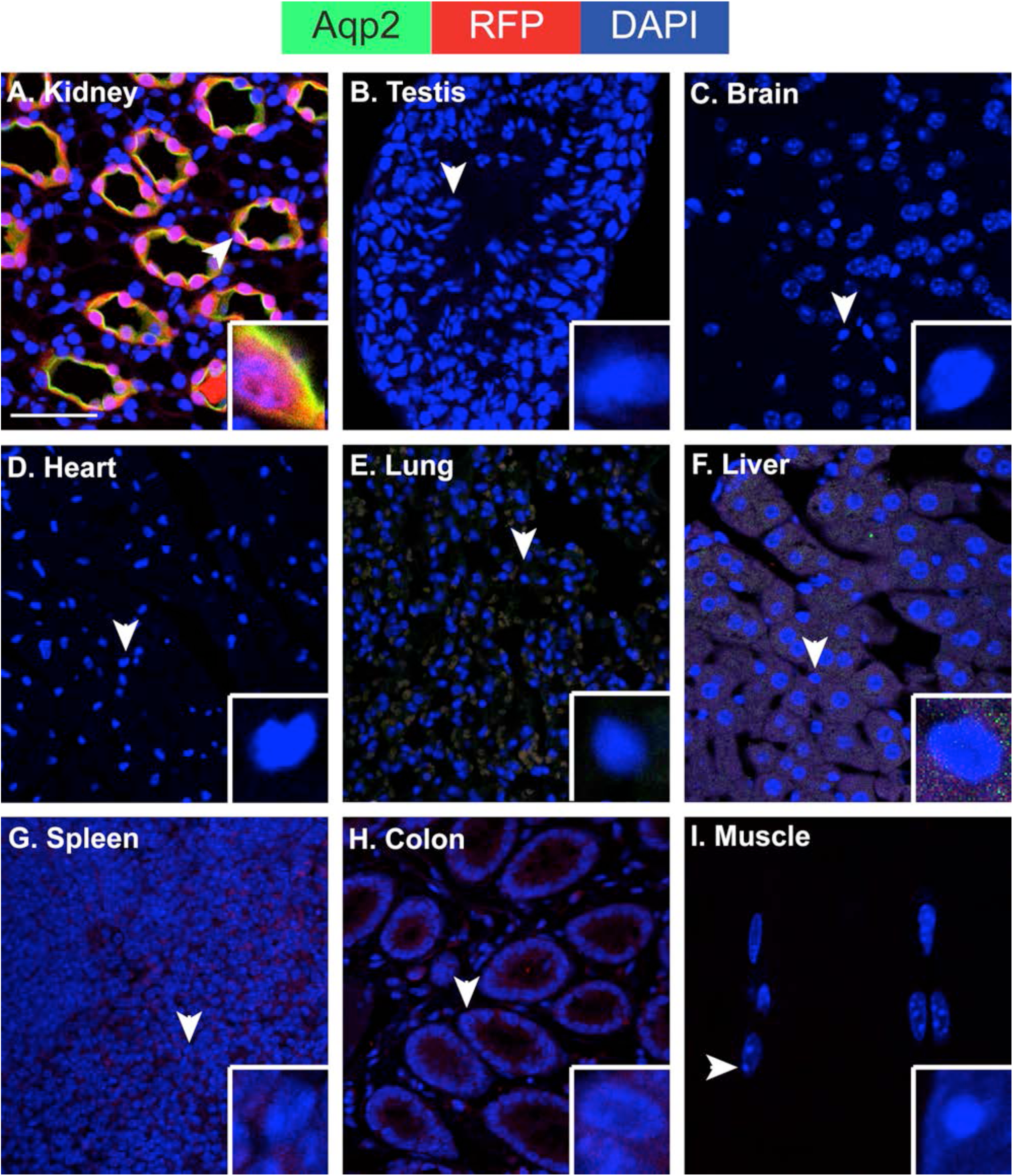
There is no indication of extrarenal Aqp2 expression and ECE-mediated recombination. *A-l.* Immunofluorescence staining for Aqp2 (green) and RFP (red) showing lack of their expression in all organs except the kidney as indicated in the adult *ECE/+ RFP/RFP* mice treated with a single dose of tamoxifen (2 mg) for 24h. Cells indicated by arrowheads were magnified in the inserts. DAPI staining (blue) for the nuclei was added. Scale bar: 50 μm and 8.3 μm for insert for each panel.

### *Aqp2^ECE^* confers high inducibility and complete fidelity in cell specificity in adult mice

To test the inducibility and fidelity, we treated two-month old *ECE/+ RFP/+* mice and *ECE/+ RFP/RFP* mice with tamoxifen (2 mg) for 24 hours. Analyses of frozen sections revealed many RFP+ cells in cortex, outer and inner medulla of both genotypes (Fig. 5 and data not shown). Immunofluorescence staining for RFP and Aqp2 showed that all RFP-labeled cells expressed Aqp2 (Fig. 6), regardless of the regions (cortex, outer and inner medulla) examined. We categorized all Aqp2^+^ cells into Aqp2^+^ RFP^+^ and Aqp2^+^ RFP^−^ cells and defined the recombination rate as the number of Aqp2^+^ RFP^+^ cells divided by all Aqp2^+^ cells in each region. For the induced *ECE/+ RFP/+* mice (n=3) the recombination rates were 46% (354/765) in the cortex, 68% (671/994) in the outer medulla and 71% (777/1093) in the inner medulla. For the induced *ECE/+ RFP/RFP* mice (n=3), the recombination rates were 46% (252/547) in the cortex, 91% (1632/1799) in the outer medulla, and 95% (1327/1401) in the inner medulla. The recombination rates in *ECE/+ RFP/+* mice were increased by either increasing the dose of tamoxifen or examination at a longer time point post induction. A single injection of tamoxifen (4 mg) increased the recombination rates to 52% (405/786), 86% (884/1026) and 90% (1062/1177) in the cortex, outer medulla and inner medulla in adult *ECE/+ RFP/+* mice 24h post injection. About 72% (2680/3728), 93% (5696/6142), and 93% (5823/6231) Aqp2^+^ cells in cortex, outer medulla, and inner medulla expressed RFP when adult *ECE/+ RFP/+* mice (n=3) were given a single injection of tamoxifen (2 mg) and analyzed 2 months later.

**Fig. 5.**
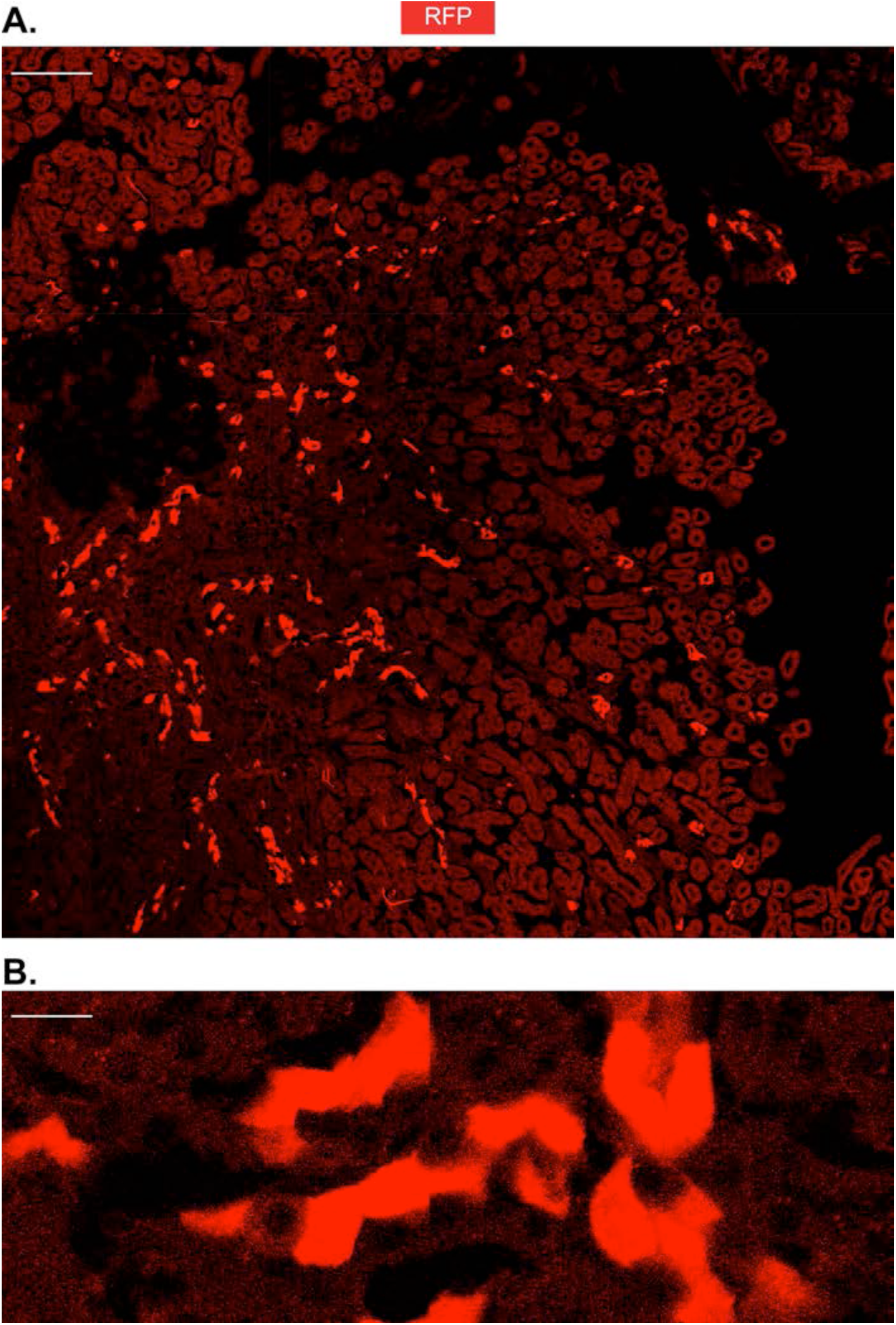
*Aqp2^ECE^* is highly inducible. A representative frozen kidney section of an adult *ECE/+ RFP/RFP* mouse treated with a single dose of tamoxifen (2 mg) for 24h showing robust RFP expression in the whole kidney. Boxed area was magnified in B. Scale bar: 200 μm for ***A*** and 20 μm for ***B.***

**Fig. 6.**
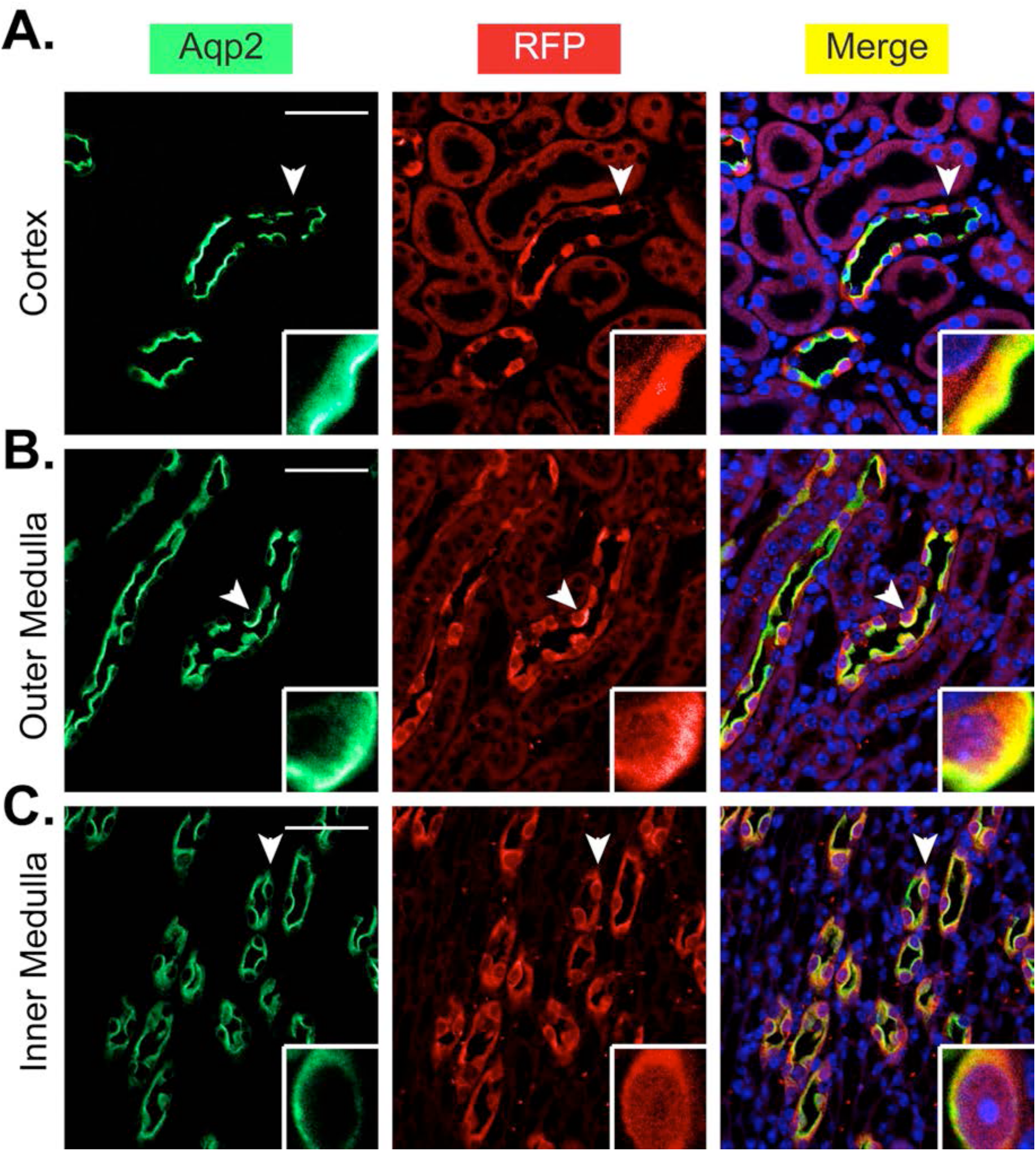
*Aqp2^ECE^* has complete fidelity in recapitulating the cell-specific expression pattern of *Aqp2. A-C*. Immunofluorescence staining for Aqp2 (green) and RFP (red) showing all RFP+ cells were also Aqp2^+^ in the whole kidney of adult *ECE/+ RFP/RFP* mice treated with a single dose of tamoxifen (2 mg) for 24h. Cells indicated by arrowheads were magnified in the inserts. Scale bar: 50 μm and 8.3 μm for insert.

To determine if RFP was co-expressed with IC markers, we first stained the kidneys of the *ECE/+ RFP/RFP* mice with three antibodies specific for RFP, Aqp2 and V-ATPase subunits B1 and B2 (B1B2) as the IC marker. We did not find a single RFP^+^ B1B2^+^ cell in the whole kidney (Fig. 7). We repeated the experiment by replacing the B1B2 antibody with an antibody specific for carbonic anhydrase II (CAII), another well-established IC marker. No RFP+ CAII+ cells were detected (Fig. 8). In brief, our data demonstrate that *Aqp2^ECE^* allele is highly inducible and mediates recombination exclusively in the Aqp2^+^ cells in adult mice.

**Fig. 7.**
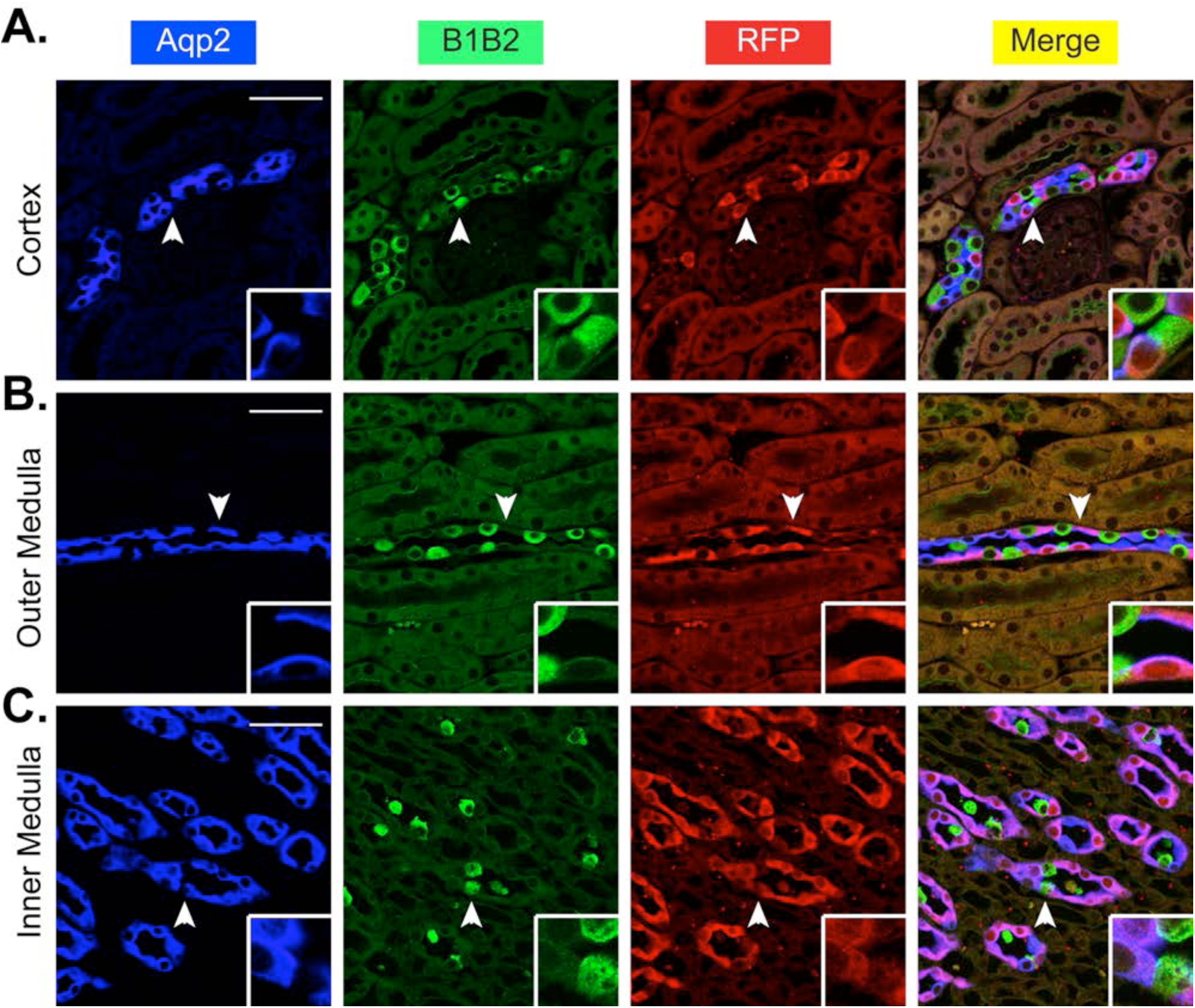
*Aqp2^ECE^* is inactive in intercalated cells marked by V-ATPase subunits B1 and B2. *A-C*. Immunofluorescence staining for Aqp2 (blue), V-ATPase B1 and B2 (B1B2, green) to mark the intercalated cells and RFP (red) showing no RFP co-expression with B1B2 in the whole kidney of adult *ECE/+ RFP/RFP* mice treated with a single dose of tamoxifen (2 mg) for 24h. Cells indicated by arrowheads were magnified in the inserts. Scale bar: 50 μm and 8.3 μm for insert.

**Fig. 8.**
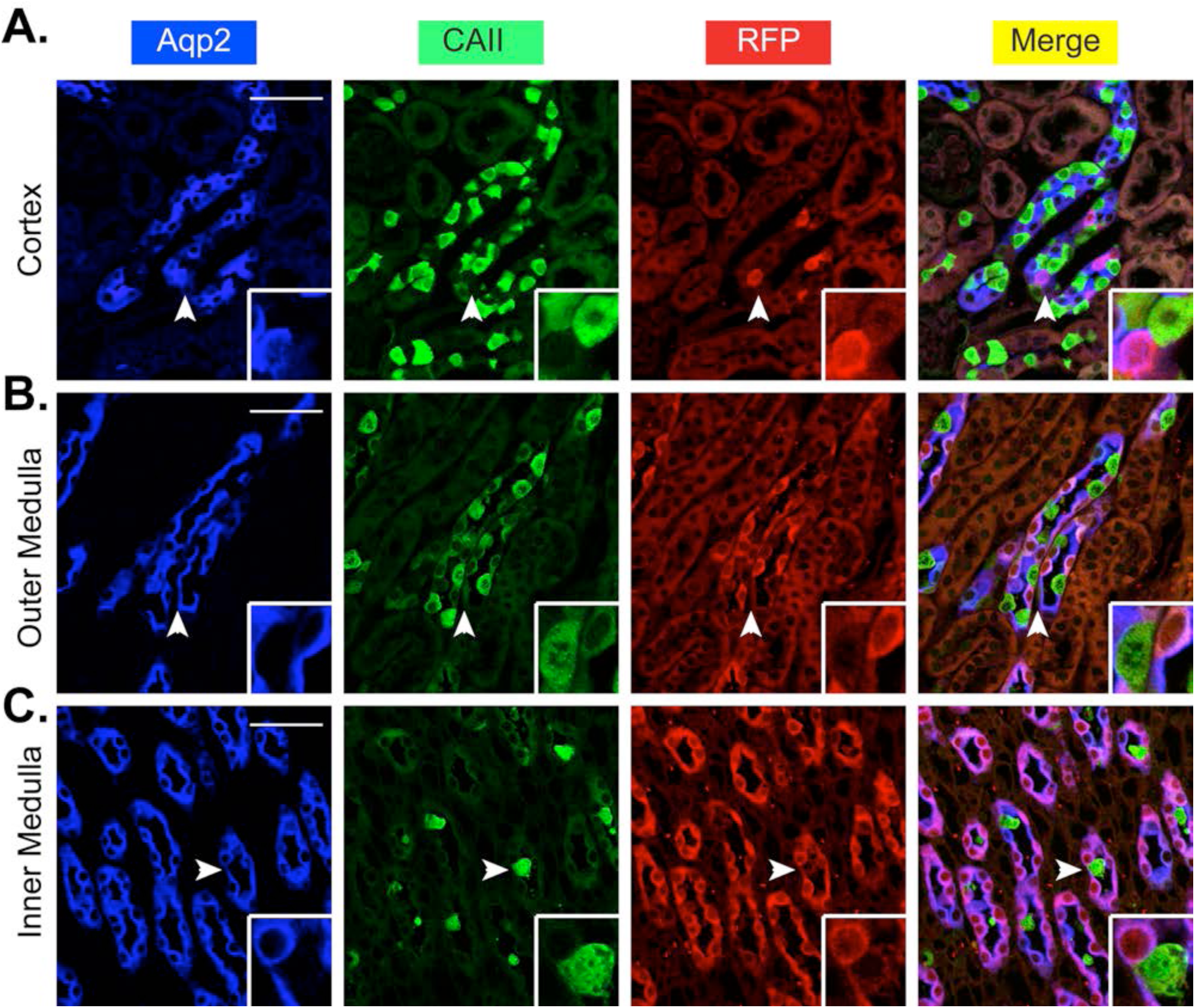
*Aqp2^ECE^* is inactive in intercalated cells marked by CAII. *A-C.* Immunofluorescence staining for Aqp2 (blue), CAII (green) to mark the intercalated cells and RFP (red) showing no RFP co-expression with CAII in the whole kidney of adult *ECE/+ RFP/RFP* mice treated with a single dose of tamoxifen (2 mg) for 24h. Cells indicated by arrowheads were magnified in the inserts. Scale bar: 50 μm and 8.3 μm for insert.

### *ECE* confers principal cell-specific recombination

In addition to Aqp2 itself, Aqp3 is also a well-established principal cell marker. To further confirm the cell specificity of ECE-mediated recombination, co-expression of RFP with Aqp3 was investigated in the kidneys of the induced *ECE/+ RFP/RFP* mice by immunofluorescence staining. As expected, all RFP^+^ cells were also positive for Aqp3 (Fig. 9).

**Fig. 9.**
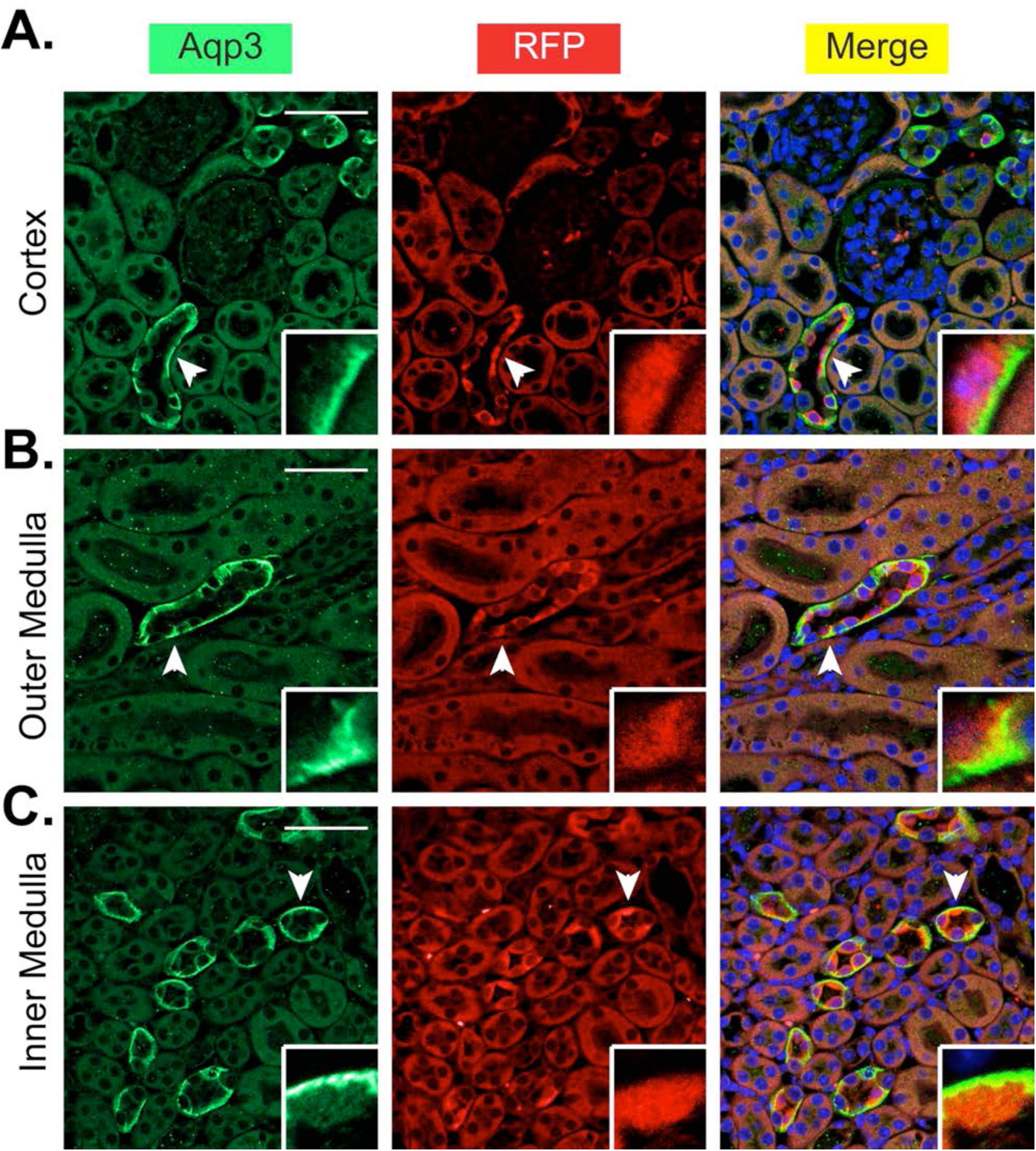
*Aqp2^ECE^* is principal-cell specific. *A-C*. Immunofluorescence staining for Aqp3 to mark the principal cells (blue) and RFP (red) showing that all RFP-expressing cells also expressed Aqp3 in the whole kidney of adult *ECE/+ RFP/RFP* mice treated with a single dose of tamoxifen (2 mg) for 24h. Cells indicated by arrowheads were magnified in the inserts. Scale bar: 50 μm and 8.3 μm for insert.

### There is no indication of extrarenal Aqp2 expression and ECE-mediated recombination

To determine if ECE-mediated recombination outside of kidney, we performed immunofluorescence staining of Aqp2 and RFP in 8 major organs (testis, brain, heart, lung, liver, spleen, colon, and muscle), with kidney as a positive control. All of these tissues were isolated from an ECE/+ RFP/RFP mice induced with tamoxifen (2 mg) for 24h. While Aqp2 and RFP were readily observed in the kidney (Fig. 10A), they were undetectable in each of all other organs tested (Fig. 10B-I).

## Discussion

To dissect the function of target genes at a particular developmental stage in a particular cell type, an ideal inducible system is required. It should strictly meet four criteria: 1) cell-specific; 2) no background recombination in the absence of induction; 3) high recombination rate after induction, and 4) complete fidelity in cell specificity to restrict recombination exclusively in cells where the endogenous gene whose promoter controls the recombinase production is expressed. Our data presented in this study demonstrate that *Aqp2^ECE^* is indeed an ideal inducible system by these criteria. Our study also for the first time offers solid evidence suggesting that the activity of ECE is absolutely dependent on tamoxifen even when ECE is expressed permanently and constantly under the control of a strong promoter such as *Aqp2* promoter, which was used in this study.

Many tamoxifen-based inducible mouse models have been developed in which *CreER^T2^* is placed under the control of a ubiquitous or a tissue/cell-specific promoter. These inducible Cre-drivers are essential reagents for temporal and spatial control of gene modifications. However, in many cases, little or no information are available with respect to leakage Cre activity in the absence of tamoxifen, recombination efficiency, and/or the faithfulness in replicating the tissue/cell-specific expression pattern of the endogenous gene whose promoter drives the CreERT^2^ recombinase. Without these data, it is a challenge for investigators to select a driver line, design experiments, and evaluate Cre recombination-induced phenotypes of their flox mice.

An argument for a much tighter control of ECE than all other three Cre fusions including CreER^T2^ has been proposed based on electroporation studies (13). Though the conclusion reached in this study agrees with our own, our case is more compelling. In our study, both the Cre-dependent RFP reporter and the *ECE* cassette are permanently inserted into the mouse genome in the defined loci. In contrast, the previous studies used electroporated plasmids encoding Cre reporters and Cre fusions. The possibility that the lack of the reporter expression simply resulted from the loss of the plasmids or the position effect of the integration site(s), although very low, cannot be completely ruled out.

Another line of evidence in support of the notion that ECE is more tightly controlled by tamoxifen than CreER^T2^ comes from the substantial background recombination of the PAC *Aqp2CreER^T2^* transgene (21). The multiple copies of the transgene, the strong *Aqp2* promoter, and possibly the integration site of the transgene result in high levels of *CreER^T2^* production. It can be speculated that the highly abundant *CreER^T2^* plus the lack of its absolute dependence on tamoxifen led to nuclear translocation of the Cre fusion to catalyze recombination in the absence of tamoxifen. Moreover, leaky activity has been reported in multiple cases including female *KspCad-CreER^T2^* (17), *Col2a1-CreER^T2^* (22), *UBC- CreER^T2^* (22) and *CAG-CreER^T2^*(14). Accumulation of the effect of even a minor leakiness over time may complicate data analyses because of the irreversibility of every recombination event. Hence, the usefulness of the leaky drivers is very limited. The limitations include that they cannot be employed to analyze the effect of disruption of a gene of interest at a particular time point and to perform in vivo lineage tracing in adult animals.

The *ECE* knock-in strategy we used in this study may be applied to other genes of interest even with strong promoters for development of various tissue/cell-specific drivers, particularly, for those having showing a leaky activity of CreER^T2^. Knock-in permits insertion of the *ECE* at a pre-defined genomic locus and ensures the presence of only a single copy of the *ECE* allele in the heterozygotes, excluding the possible overproduction of the Cre fusion due to an increased copy number.

Three mechanisms can be speculated for a tighter control of the ECE than CreER^T2^ (13). Compared to CreER^T2^, ECE may form a tighter inactive complex with Hsp90 in the cytoplasm, remain inactive even after losing one ER domain, and be less active. However, the high recombination rate of *Aqp2^ECE^* after a single dose of tamoxifen administration argues against the less activity of ECE.

One injection of tamoxifen was sufficient to induce robust RFP expression after only 24h in *ECE/+ RFP/+* and *ECE/+ RFP/RFP* mice. The recombination efficiency can reach up to 95%. Many Cre drivers exhibited substantially lower recombination rates even after 3-5 daily injections of tamoxifen. For example, *KspCad-CreER^T2^* yielded a recombination efficiency of only 40-50% even after administration of tamoxifen at a much higher dose (5 X 5 mg/day) (11). *SLC34a1GCE* induced with five doses of 3 mg of tamoxifen mediated 58% recombination in cortical S3 segment (10). Injection of tamoxifen in adult *Podocin-iCreER^T2^* -containing mice once resulted in only 10% excision (23).

Since the *ECE* was inserted at the ATG of the endogenous *Aqp2* locus, it is under the control of the same regulatory elements that governs Aqp2 expression. As expected, RFP signals as the output of the ECE-mediated recombination recapitulated the principal cell-specific expression of the endogenous *Aqp2* in the CD with complete fidelity in cell specificity, although some Aqp2^+^ cells were RFP^−^. We verified the fidelity with a high resolution at single cells by co-staining Aqp2 and RFP with or without IC markers on the same slides. A previous report using RT-PCR and immunohistochemistry demonstrated Aqp2 expression in the male reproductive system of C57BL/CBA mice (15). However, we did not detect Aqp2 and thus RFP expression in the testis of the induced *ECE/+ RFP/RFP* mice with highly pure C57BL/6 background. Whether this inconsistency stems from the difference in the genetic background of the mouse models used or something else requires further investigation.

The cell-specific property distinguishes the *Aqp2^ECE/+^* from the vast majority of the existing kidney-specific inducible mouse models, which are not cell-specific. For instance, *KspCad-CreER^T2^* transgene activates *R26R-EYFP* reporter in proximal tubules, thick ascending limbs of loops of Henle, distal convoluted tubule, and collecting duct (17). The sodium-dependent inorganic phosphate transporter *SLC34a1* and thus *SLC34a1-GFPCreER^T2^* are expressed in differentiated proximal tubule, which consists of S1, S2 and S3 segments (10). Protrudin, which is encoded by *Zfyve27,* is expressed in interstitial cells, collecting duct cells, and thin limbs of Henle’s loop in the kidney papilla (16). Faithful recapitulation of Protrudin is expected to confer *Zfyve27-CreER^T2^* being active in these cell types. Hence, *Aqp2^ECE/+^* mice should be very powerful and complementary to the increasing list of inducible mouse model systems for renal research.

The most obvious application of the *Aqp2^ECE/+^* mice is to inactivate floxed targeted genes temporally and specifically in the principal cells in CD and presumably in CNT. A typical example is deletion of *Pkd1* and *Pkd2* in adult mice. Like many other disorders such as various types of cancer, polycystic kidney disease (PKD) is a focal disease in nature and involves somatic mutations of target genes. In this adult-onset disease cysts develop in <1% of the nephrons. While renal cysts in autosomal dominant polycystic kidney disease can arise from cells throughout the nephron (18), most cysts stain positive for markers of collecting duct cells in advanced autosomal dominant polycystic kidney disease kidneys. Therefore, CD/CNT-specific inactivation of *Pkd1* and *Pkd2* in adult mice using *Aqp2^ECE/+^* may offer critical insights into the molecular and pathological basis of PKD.

*Aqp2^ECE/+^* can also be employed to temporally activate genes of interest by removing a “STOP” cassette that prevents their expression within the CD. This is clearly demonstrated by the expression of RFP in this study. Such applications will undoubtedly enhance our ability to genetically dissect the role of putative disease-causing genes within either the developing or adult kidney.

Given the absolute no leakiness and complete fidelity in cell specificity, *Aqp2^ECE/+^* mice will provide a unique advantage to trace Aqp2^+^ lineage. With the abolished histone H3 K79 dimethylation as a tracing marker, we have showed that the principal cells and most intercalated cells are derived from a new pool of progenitor cells expressing Aqp2 in *Dot1l^AC^* mice during development (25). However, whether Aqp2^+^ progenitor cells exist in adult kidney remains completely unknown. *Aqp2^ECE/+^* mice will become a unique tool to address this question.

Like all other constitutive or inducible Cre drivers, *Aqp2^ECE^* did not yield a 100% recombination rate under three conditions tested for *ECE/+ RFP/+* mice (2 and 4 mg tamoxifen for 24h, and 2 mg tamoxifen for 2 months) and one condition for *ECE/+ RFP/RFP* mice (2 mg tamoxifen for 24h). The recombination rates in the cortex, outer and inner medulla may be further increased via a variety of methods including repeated tamoxifen injections, higher doses of tamoxifen, examination at even longer time points post injection, and water deprivation or restriction to increase the *Aqp2* promoter activity. It is also possible that the rates may not be significantly changed due to very high recombination rates we have already observed, the potential intrinsic limitation of the *Aqp2^ECE^* model system, or both. Nevertheless, these possibilities have not been completely addressed, a limitation of the current study. *Aqp2^ECE^* can presumably be used for genetic modifications in the CNT. However, immunofluorescence staining experiments with CNT-specific markers are required to conclusively demonstrate ECE-mediated recombination in the CNT.

## Acknowledgment

We thank the UCI Transgenic Mouse Facility (TMF) for ES targeting and production of chimeric mice. The UCI TMF is a shared resource funded in part by the Chao Family Comprehensive Cancer Center Support Grant (P30CA062203) from the National Cancer Institute. This study was started when Dr. Lihe Chen pursued his Ph.D. in Dr. Zhang’s lab in the University of Texas Health Science Center at Houston.

## List of nonstandard abbreviations

CNT/CD: connecting tubule/collecting duct
GFR: glomerular filtration rate
ER: estrogen receptor
ECE: ER^T2^CreER^T2^
4OHT: 4-hydroxytamoxifen
RFP: red fluorescent protein
LBD: ligand-binding domain
BP: blood pressure
CAII: Carbonic anhydrase II

